# DoReMiTra: An R/Bioconductor data package for orchestrating the analysis of radiation transcriptomic studies

**DOI:** 10.64898/2025.12.12.691104

**Authors:** Ahmed Salah, Sebastain Zahnreich, Federico Marini

## Abstract

**Summary:** Understanding the molecular impact of ionizing radiation exposure is essential for both biomedical research and public health. Among the possible approaches to study this phenomenon, gene expression profiling via transcriptomics assays has been a valuable approach over the last decades to unravel the mechanisms of cellular responses to radiation. To our knowledge, there is no data package gathering well-curated radiation transcriptomic datasets covering microarrays and, more recently, RNA sequencing.

Therefore, we present DoReMiTra, an R/Bioconductor data package that represents the first unified radiation transcriptomics dataset collection integrated with Bioconductor’s ExperimentHub for efficient distribution.

DoReMiTra standardizes and harmonizes sample-level metadata and provides pre-processed SummarizedExperiment (SE) objects to facilitate comparative analyses. Additionally, we introduce a lightweight Shiny app interface for interactive visualization and preliminary exploration. DoReMiTra serves as a valuable resource and tool in radiation research for benchmarking, integrative analyses, and biomarker discovery.

**Availability and Implementation:** DoReMiTra is available under the MIT license at https://bioconductor.org/packages/DoReMiTra.

## Introduction

Ionizing radiation (IR) is a long-standing cornerstone of medical practice, essential for diagnostic imaging and the treatment of both benign and malignant diseases. However, despite its significant clinical value, planned and especially accidental IR exposures still pose health risks (Ron 2002).

Notably, radiation biology research increasingly relies on transcriptomic profiling to uncover the molecular mechanisms underlying cellular responses to (Kabacik *et al*. 2011; Manning *et al*. 2017) IR. However, the diversity of radiation qualities, each with distinct energy deposition patterns and biological effects, can lead to diverse transcriptomic responses depending on dose, dose-rate, radiation type, and biological context (Ismail, Hamad and Harki 2012; Michalettou *et al*. 2021; Talapko *et al*. 2024). Therefore, gene expression profiling has emerged as a rapid and sensitive tool for dose estimation and toxicity prediction in clinical, biodosimetry, and triage applications (Dressman *et al*. 2007; Meadows *et al*. 2008).

Numerous gene expression studies have investigated these effects across human and animal models (El-Saghire *et al*. 2013; Hall *et al*. 2017). Still, data reuse is hindered by inconsistent metadata annotation, variable preprocessing methods, and a lack of accessibility, which makes it challenging to define experimental factors for statistical analysis (Koesten *et al*. 2020; Bustos *et al*. 2025). To date, no resource integrates radiation transcriptomics datasets into a unified structure. To address this, we present DoReMiTra (radiation DOse REsponse Measured In TRAnscriptomics), the first and comprehensive data package that aggregates blood transcriptomic datasets from experiments involving photons (X-rays, ***γ***-rays) and neutrons, providing access to well-documented and preprocessed transcriptomic datasets. We focused mainly on datasets generated from exposed blood samples since it is a very radiosensitive tissue that is always exposed to radiation in both medical and accidental exposure scenarios (Rothkamm *et al*. 2013; Heylmann *et al*. 2014).

The package includes datasets primarily retrieved from GEO via GEOquery (Davis and Meltzer 2007), unified under a common structure, with metadata covering radiation type, dose, time point, sex, organism, and experiment setting (e.g., in vivo, ex vivo). This data package not only streamlines access to a curated radiation transcriptomic landscape but also all the datasets included are provided as SE objects that can be easily sourced from the Bioconductor ExperimentHub ecosystem (Morgan and Shepherd 2025) to allow seamless integration with R/Bioconductor workflows (Robinson, McCarthy and Smyth 2010; Love, Huber and Anders 2014; Ritchie *et al*. 2015; Ludt *et al*. 2022), simplifying the comparability across studies and supporting reproducible analysis pipelines. This integration would make it feasible for biologists, bioinformaticians, and statisticians to conduct the analysis and integration without the hassle of metadata inconsistency across studies. DoReMiTra, accompanied by a Shiny app for exploratory visualization, serves as a valuable resource in radiation research and a tool for developing biomarkers of radiation exposure in the context of radiation therapy, radiation oncology, and biodosimetry, and encourages reproducibility and data reuse in radiation transcriptomic research (Wilkinson *et al*. 2016; Fraga-González *et al*. 2025).

**Figure 1.**
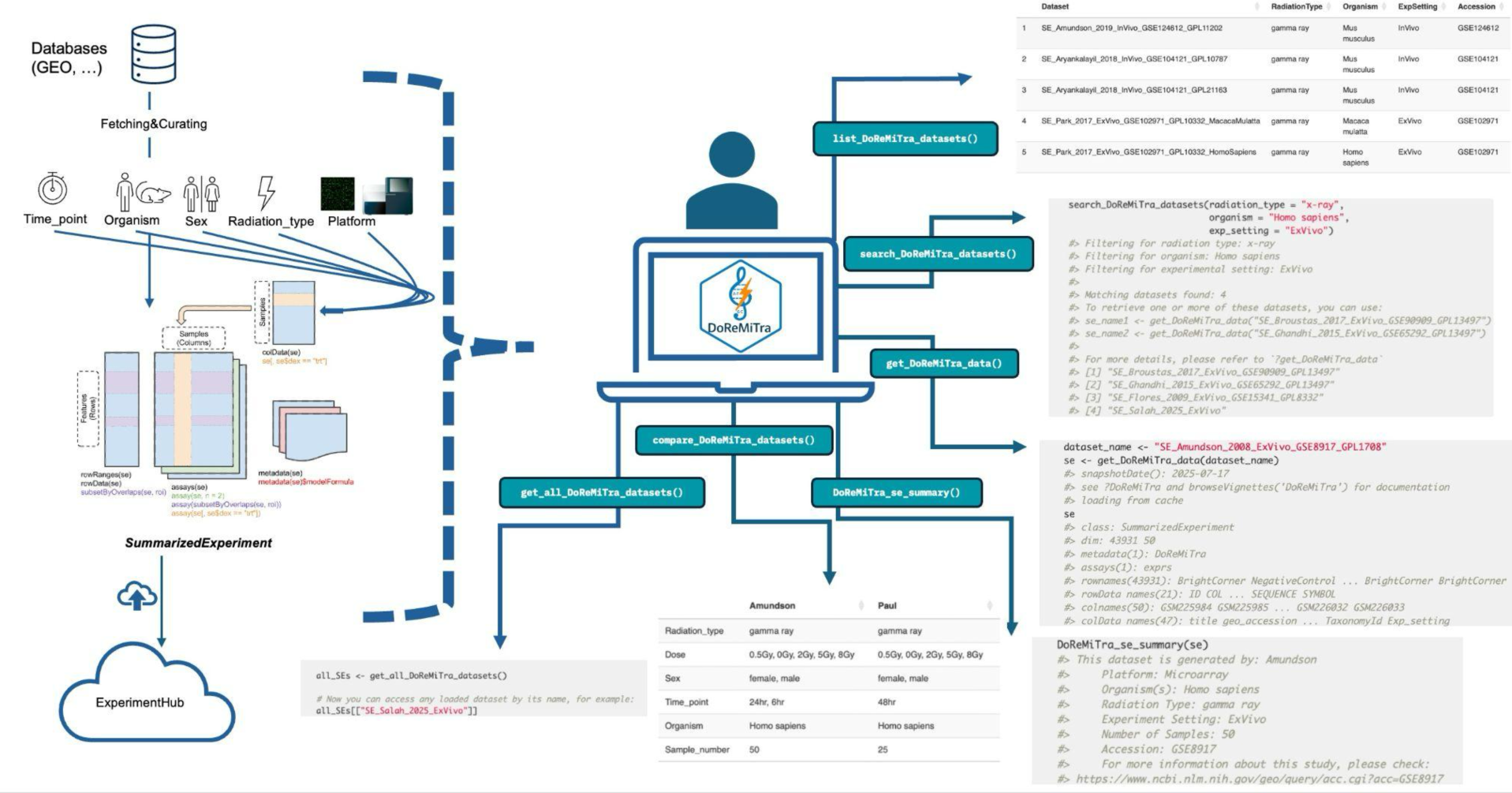
Graphical representation of the creation and typical usage workflow for DoReMiTra. The package provides: get_DoReMiTra_data() retrieves a dataset by name. list_DoReMiTra_datasets() views metadata for all available datasets. search_DoReMiTra_datasets() filters datasets by specified metadata fields. compare_DoReMiTra_datasets() compares metadata fields across datasets. DoReMiTra_se_summary() displays a summary of key metadata and GEO links. get_all_DoReMiTra_datasets() retrieves all datasets from the DoReMiTra collection.

### Data Collection and Processing

Many publicly available radiation transcriptomic datasets contain non-standardized or inconsistent metadata. In some studies, even samples from the same experiment are annotated with heterogeneous terminology or are described by overly long, unstructured text strings. These inconsistencies hinder the reliable extraction of key experimental factors such as dose, time point, or sex, and they complicate the construction of appropriate statistical design formulas for downstream analyses in packages such as limma, DESeq2, or edgeR.

To address this, the first building block of our package build was collecting datasets from irradiated blood samples of human or mouse, primarily from GEO, based on relevance to biodosimetry. Priority was given to experiments involving X-rays, ***γ***-rays, and neutrons, with well-defined dose and time point information. The GEO queries to obtain all the datasets are available on the following link (https://github.com/AhmedSAHassan/DoReMiTra/blob/devel/inst/extdata/Geo_Queries.qmd).

All the information about each dataset, including accession number, author name, type of radiation, dose, experimental setting, etc., was collected in a tabular format. The datasets were then programmatically fetched by their accession number and processed into SE objects using standardized pipelines. Sample-level and feature-level metadata, reported in the respective colData and rowData slots, were manually curated with particular focus on adopting unified terminologies across all the datasets, including Platform, Radiation_type, Dose, Organism, Time_point, Sex, and Exp_setting, to facilitate the statistical analysis and integration. When metadata fields were missing or inconsistently reported, information was cross-validated using associated publications.

The RNA-seq dataset is provided as raw counts, whereas the normalization methods for the microarray datasets vary across studies, according to the decisions made by the original investigators due to platform-specific differences. Accordingly, a link to each dataset on GEO is included in the metadata of the corresponding SE object.

The current release includes 36 datasets, providing a high number of samples for integrative analysis. For example, the package provides 14 datasets for *ex vivo* ***γ***-irradiated human blood samples with a total of 753 samples included. More datasets will be regularly included in the collection as they are generated and made publicly available. The current collection of datasets covers: Organisms (*Homo sapiens, Mus musculus, Macaca mulatta)*, radiation types (X-ray, gamma ray, neutron), experimental settings (in vivo, ex vivo), time points (minutes to days) post-exposure, sex (male and female), and dose (Gy)

A detailed processing pipeline can be found in the Rmarkdown file “make-data-DoReMiTra.Rmd” from the installed “scripts” folder of the DoReMiTra package.

### The DoReMiTra workflow

The DoReMiTra functionality includes a series of steps. The package built and workflow are shown in Figure1. The primary entry point for exploring the package is list_DoReMiTra_datasets(), which returns a metadata data frame summarizing all datasets available in the DoReMiTra collection, including key attributes such as title, organism, radiation type, experimental setting, and accession numbers. Secondly, get_all_DoReMiTra_Datasets() retrieves all datasets from ExperimentHub and returns them as a named list of SummarizedExperiment (SE) objects. This function is particularly useful for batch processing or for global exploration of all curated radiation transcriptomic datasets. File retrieval is handled through BiocFileCache, which enables efficient caching of files.

Users can refine their selection using search_DoReMiTra_datasets(), which filters available datasets based on metadata fields such as radiation type, organism, or experimental setting, allowing rapid identification of datasets relevant to a specific research question.

Once a dataset of interest is selected, get_DoReMiTra_data() fetches the corresponding SE object from ExperimentHub within seconds. The function includes a gene_symbol argument, which, when set to TRUE, sets rownames to gene symbols when available. In cases of duplicated symbols, the gene symbol and probe ID are appended; if the gene symbol is missing (NA), the probe ID is used instead. Setting gene_symbol = FALSE retains probe IDs as rownames. Once the SE object is retrieved by this function, it can be directly explored by DoReMiTra_explorer(), an interactive Shiny application for visualization and exploratory analysis of the DoReMiTra datasets. The app enables users to intuitively explore transcriptomic responses to radiation through a suite of customizable plots, including principal component analysis (PCA) with user-defined gene subsets and color schemes based on experimental factors, boxplots for expression distributions, dose–response gene plots, and heatmaps of highly variable genes. Users can adjust parameters such as the number of genes displayed and the inclusion of clustering, offering flexibility for both overview and detailed inspection of radiation-induced transcriptional patterns.

The Shiny app can be found in the GitHub repository (https://github.com/AhmedSAHassan/DoReMiTraExplorer).

A quick overview of any retrieved dataset can be obtained using summarize_DoReMiTra_se(), which reports the number of samples, available metadata fields, and key experimental characteristics such as radiation type, organism, and platform. Finally, **compare_DoReMiTra_datasets()** allows users to compare two or more SE objects side-by-side by summarizing essential metadata attributes, including radiation type, dose, time point, and experimental settings.

An exemplary integrative analysis of datasets retrieved via DoReMiTra is presented in Supplementary File 1, with the source code for the notebook provided in the GitHub repository https://github.com/AhmedSAHassan/DoReMiTraSup.

## Discussion and future directions

DoReMiTra provides a one-stop shop of harmonized and accessible datasets for blood transcriptomic studies investigating the biological effects of various radiation types, including X-rays, ***γ***-rays, and neutrons. By curating publicly available GEO datasets into standardized SE objects and embedding them within Bioconductor’s ExperimentHub, DoReMiTra enables seamless access, reproducibility, and interoperability with the broader Bioconductor ecosystem. This resource bridges a critical gap in radiation biology by offering a standardized structure and harmonized metadata; it lays the groundwork for integrative analyses, reproducible benchmarking, and the identification of radiation-responsive molecular signatures.

Although DoReMiTra does not embed a full framework for statistical analysis of the provided datasets, DoReMiTra substantially lowers the barrier to performing radiation transcriptomics research by providing the largest curated collection of harmonized datasets in an analysis-ready format. We refer the reader to the analysis of Salah et al (Salah *et al*. 2025), which has been described in detail in the repository (https://github.com/AhmedSAHassan/Phybion_ShortReads_Analysis), which also illustrates how proper design specification is essential to extract robust radiation signatures, especially when working with human data.

Future developments of DoReMiTra will include modules for cross-platform normalization, which would be particularly valuable for meta-analysis and machine learning workflows that require comparable expression scales across datasets. Similarly, batch correction is intentionally left to downstream analyses, allowing users to apply appropriate methods (e.g., ComBat, RUVSeq, sva) (Leek *et al*. 2012; Risso *et al*. 2014; Zhang, Parmigiani and Johnson 2020). This modular approach maintains flexibility, enabling researchers to tailor normalization and integration strategies to their analytical needs while ensuring reproducibility and methodological transparency. Additionally, we plan to include dataset-specific gene signatures, which will further facilitate cross-study comparisons and the identification of conserved radiation-responsive transcriptional patterns across different exposure types and experimental systems. As the current version includes datasets from irradiated blood samples due to their biological relevance, future updates will include other tissue types to broaden the spectrum to answer different research questions.

Beyond facilitating immediate reuse of existing data, DoReMiTra encourages the community to adopt transparent and interoperable data practices, ultimately strengthening research in radiobiology, biodosimetry, and radiotherapy. Additionally, DoReMiTra can provide valuable resources for training and education due to its diverse collection of well-structured datasets for multiple comparisons. The package promotes FAIR data principles by being findable (ExperimentHub search), accessible (programmatic retrieval), interoperable (adopting Bioconductor SE structure), and reusable (curated metadata and documentation). Taken together, DoReMiTra offers a solution for accessing, exploring, and analyzing curated radiation transcriptomic datasets, enabling researchers to perform integrative studies and accelerate the discovery of radiation-responsive biomarkers.

## Data and code availability

The DoReMiTra package is available on Bioconductor under the MIT license at https://bioconductor.org/packages/DoReMiTra; the companion web application to explore the included datasets is available at https://github.com/AhmedSAHassan/DoReMiTraExplorer. The source code for the analysis notebook presented in the Supplementary File is available in the GitHub repository https://github.com/AhmedSAHassan/DoReMiTraSup.

## Competing interests

No competing interest is declared.

## Author contributions statement

A.S. Data curation, methodology, software, validation, writing – original draft, writing – review & editing S.Z. Supervision, funding acquisition, project administration, resources, writing – review & editing

F.M. Conceptualization, methodology, software, supervision, funding acquisition, project administration, resources, writing – original draft, writing – review & editing

## Acknowledgments

This study was supported by the German Federal Ministry of Education and Research, Grant 02NUK084A (A.S.). The work of FM is also supported by the Deutsche Forschungsgemeinschaft (DFG, German Research Foundation) Projektnummer 318346496 - SFB1292/2 TP19N.

## Notes

### Competing Interest Statement

The authors have declared no competing interest.

