## Supplementary File 1 for "DoReMiTra: An R/Bioconductor data package for orchestrating the analysis of radiation transcriptomic studies"

### Using DoReMiTra for analyzing and integrating publicly available transcriptomics radiation datasets

AUTHORS | AFFILIATIONS
Ahmed Salah — Department of Radiation Oncology and Radiation Therapy, Mainz, Germany; Institute of Medical Biostatistics, Epidemiology and Informatics (IMBEI), Mainz, Germany
Sebastian Zahnreich — Department of Radiation Oncology and Radiation Therapy, Mainz, Germany
Federico Marini — Institute of Medical Biostatistics, Epidemiology and Informatics (IMBEI), Mainz, Germany

PUBLISHED
December 12, 2025

#### 1 Loading the required packages

Here we load all packages required to reproduce the analysis in this notebook.

```
library("DoReMiTra")
library("mosdef")
library("org.Hs.eg.db")
library("DESeq2")
library("ggplot2")
library("ComplexHeatmap")
library("VennDiagram")
library("limma")
library("circlize")
library("GeneTonic")
library("clusterProfiler")
library("patchwork")
```

#### 2 Searching for datasets

The user can use `search_DoReMiTra_datasets()` to filter the available DoReMiTra datasets based on metadata fields such as radiation type, organism, or experimental setting. This function helps narrow down datasets of interest before fetching them.

```
# X-ray datasets
search_DoReMiTra_datasets(radiation_type = "X-ray",
                          organism = "Homo sapiens",
                          exp_setting = "ExVivo")
#> Filtering for radiation type: X-ray
#> Filtering for organism: Homo sapiens
#> Filtering for experimental setting: ExVivo
#>
#> Matching datasets found: 4
#> To retrieve one or more of these datasets, you can use:
#> se_name1 <- get_DoReMiTra_data("SE_Broustas_2017_ExVivo_GSE90909_GPL13497")
#> se_name2 <- get_DoReMiTra_data("SE_Ghandhi_2015_ExVivo_GSE65292_GPL13497")
#>
#> For more details, please refer to `?get_DoReMiTra_data`
#> [1] "SE_Broustas_2017_ExVivo_GSE90909_GPL13497"
#> [2] "SE_Ghandhi_2015_ExVivo_GSE65292_GPL13497"
#> [3] "SE_Flores_2009_ExVivo_GSE15341_GPL8332"
#> [4] "SE_Salah_2025_ExVivo"

# gamma ray datasets
search_DoReMiTra_datasets(radiation_type = "gamma ray",
                          organism = "Homo sapiens",
                          exp_setting = "ExVivo")
#> Filtering for radiation type: gamma ray
#> Filtering for organism: Homo sapiens
#> Filtering for experimental setting: ExVivo
#>
#> Matching datasets found: 14
#> To retrieve one or more of these datasets, you can use:
#> se_name1 <- get_DoReMiTra_data("SE_Park_2017_ExVivo_GSE102971_GPL10332_HomoSapiens")
```

```
#> se_name2 <- get_DoReMiTra_data("SE_Vasilyev_2017_ExVivo_GSE97000_GPL17077")
#>
#> For more details, please refer to `?get_DoReMiTra_data`
#>  [1] "SE_Park_2017_ExVivo_GSE102971_GPL10332_HomoSapiens"
#>  [2] "SE_Vasilyev_2017_ExVivo_GSE97000_GPL17077"
#>  [3] "SE_Paul_2013_ExVivo_GSE44201_GPL6480"
#>  [4] "SE_Paul_2013_ExVivo_GSE44201_GPL6848"
#>  [5] "SE_Nosel_2013_ExVivo_GSE43151_GPL13497"
#>  [6] "SE_Girardi_2012_ExVivo_GSE20173_GPL6480"
#>  [7] "SE_Amundson_2011_ExVivo_GSE23515_GPL6480"
#>  [8] "SE_Gruel_2008_ExVivo_GSE6978_GPL4803"
#>  [9] "SE_Rouchka_2019_ExVivo_GSE63952_GPL15207"
#> [10] "SE_Amundson_2008_ExVivo_GSE8917_GPL1708"
#> [11] "SE_Manikandan_2014_ExVivo_GSE36355_GPL6883"
#> [12] "SE_Lee_2013_ExVivo_GSE44245_GPL570"
#> [13] "SE_Rouchka_2015_ExVivo_GSE64375_GPL6244"
#> [14] "SE_Ankermit_2015_ExVivo_GSE55953_GPL14550"
```

### 3 Retrieving the datasets

After selecting the dataset of interest, `get_DoReMiTra_data()` can be used to fetch a `SummarizedExperiment` (SE) object directly from ExperimentHub.

```
# we start with a recent study by Salah et al., conducted on RNA-seq
dataset_name_1 <- "SE_Salah_2025_ExVivo"

# we would like also to compare it to another dataset from another radiation type to see share
dataset_name_2 <- "SE_Paul_2013_ExVivo_GSE44201_GPL6848"

se_salah <- get_DoReMiTra_data(dataset_name_1)
#> see ?DoReMiTra and browseVignettes('DoReMiTra') for documentation
#> loading from cache
se_paul <- get_DoReMiTra_data(dataset_name_2, gene_symbol = TRUE)
#> see ?DoReMiTra and browseVignettes('DoReMiTra') for documentation
#> loading from cache
```

These simple commands provide the user an analysis-ready object for the datasets generated by (Salah et al. 2025) and by (Paul, Smilenov, and Amundson 2013).

### 4 Obtaining summaries of the datasets

To quickly obtain a summary of an SE object, use `summarize_DoReMiTra_se()`, which provides a concise overview including the number of samples, genes, radiation types, and other metadata.

```
summarize_DoReMiTra_se(se_salah)
#> This dataset is generated by: Salah
#>     Platform: RNAseq
#>     Organism(s): Homo sapiens
#>     Radiation Type: X-ray
#>     Experiment Setting: ExVivo
#>     Number of Samples: 30
#>     Accession: NA
#>     For more information about this study, please check:
#> https://github.com/AhmedSAHassan/Phybion_ShortReads_Analysis
summarize_DoReMiTra_se(se_paul)
#> This dataset is generated by: Paul
#>     Platform: Microarray
#>     Organism(s): Homo sapiens
#>     Radiation Type: gamma ray
#>     Experiment Setting: ExVivo
#>     Number of Samples: 50
#>     Accession: GSE44201
#>     For more information about this study, please check:
#> https://www.ncbi.nlm.nih.gov/geo/query/acc.cgi?acc=GSE44201
```

Once we retrieved the analysis-ready datasets of intereset, we can explore them and analyse them in detail.

### 5 Comparing the metadata of multiple datasets

To compare key metadata information from two or more SE objects from the DoReMiTra collection, `compare_DoReMiTra_datasets()` summarizes key metadata information such as radiation type, dose, and time point.

```
se_list<- list(
  `Salah et al.` = se_salah,
  `Paul et al.` = se_paul
)
compare_DoReMiTra_datasets(se_list = se_list) |>
  kableExtra::kable()
```

| | Salah et al. | Paul et al. |
|---|---|---|
| Radiation_type | X-ray | gamma ray |
| Dose | 0.5Gy, 0Gy, 1Gy, 2Gy, 4Gy | 0.5Gy, 0Gy, 2Gy, 5Gy, 8Gy |
| Sex | Female, Male | female, male |
| Time_point | 2hrs, 6hrs | 24hr, 6hr |
| Organism | Homo sapiens | Homo sapiens |
| Sample_number | 30 | 50 |

### 6 Using DoReMiTraExplorer for exploratory data analyses

One can also explore the datasets with `DoReMiTraExplorer`, a companion app package we have developed.

The command to install the app is exemplified in the following chunk:

```
# Install the DoReMiTra-explorer Shiny App
devtools::install_github("AhmedSAHassan/DoReMiTraExplorer")
```

The retrieved SummarizedExperiment objects can be explored with `DoReMiTra_explorer()`, an interactive Shiny application for visualization and exploratory analysis of DoReMiTra datasets. This app allows users to examine transcriptomic responses to radiation using:

- Principal Component Analysis (PCA)
- Boxplots for the distribution of expression values
- Dose-response gene plots
- Heatmaps of highly variable genes

It provides flexibility for both high-level overviews and detailed inspection of radiation-induced transcriptional patterns.

To load the app, we can simply call

```
# load the DoReMiTraExplorer package
library("DoReMiTraExplorer")

# launch the app if in an interactive session
if(interactive()) {
  DoReMiTraExplorer::DoReMiTra_explorer(se = se_salah)
}
```

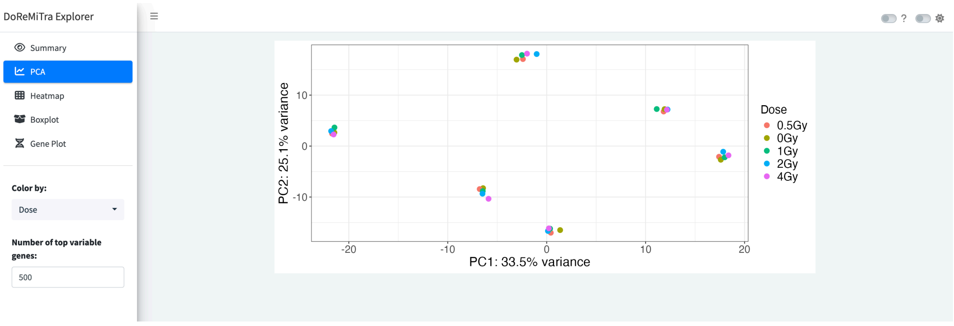


Snapshot of the DoReMiTraExplorer app

### 7 Analyzing the datasets, in detail

Once we identified and retrieved the analysis-ready datasets of intereset, we can explore and analyse them.

#### 7.1 Working on the X-ray dataset

As we could observe from the PCA plot obtained via `DoReMiTraExplorer`, the donor and time effects are the main sources of variability. These factors should be accounted for in downstream statistical analyses. Specifically, we use in this case the DESeq2 statistical framework, and focus on the **2Gy vs 0Gy** comparison, starting from the subset of (Salah et al. 2025) including samples 2 hours after irradiation.

This dataset consists of 3 donors, 5 doses (0, 0.5, 1, 2, and 4Gy), and 2 timepoints (2h and 6h)

```
dds_salah <- DESeqDataSet(se_salah, design = ~1)
#> using counts and average transcript lengths from tximeta

dds_salah$Dose <- factor(dds_salah$Dose)
dds_salah$Donors <- factor(dds_salah$Donors)
dds_salah$Dose <- relevel(dds_salah$Dose, ref = "0Gy")

dds_salah_2h <- dds_salah[, dds_salah$Time_point == "2hrs"]
dds_salah_6h <- dds_salah[, dds_salah$Time_point == "6hrs"]

vsd_salah <- vst(dds_salah)
#> using 'avgTxLength' from assays(dds), correcting for library size

pca_plot <- plotPCA(vsd_salah,
                    intgroup = c("Donors", "Time_point"),
                    returnData = TRUE)
#> using ntop=500 top features by variance

percentVar <- round(100 * attr(pca_plot, "percentVar"))

PCA <- ggplot(pca_plot, aes(x = PC1, y = PC2, color = Donors, shape = Time_point)) +
  geom_point(size = 3) +
  scale_shape_manual(values = c(16, 17)) +
  xlab(paste0("PC1: ", percentVar[1], "% variance")) +
  ylab(paste0("PC2: ", percentVar[2], "% variance")) +
  theme_minimal()+
  theme(plot.background = element_rect(fill = "white", color = NA))

PCA
```

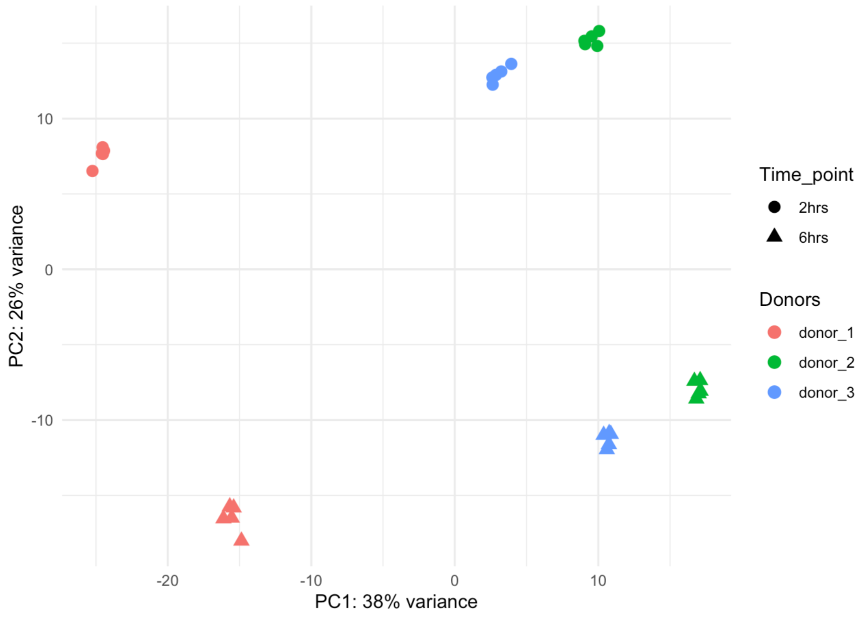


```
vsd_salah_6h <- vst(dds_salah_6h)
#> using 'avgTxLength' from assays(dds), correcting for library size

pca_donors <- plotPCA(vsd_salah_6h,
        intgroup = "Donors",
        returnData = FALSE) +
  theme_minimal() +
  labs(title = "Salah et al., PCA by donors")
#> using ntop=500 top features by variance
pca_dose <- plotPCA(vsd_salah_6h,
        intgroup = "Dose",
        returnData = FALSE) +
  theme_minimal() +
  labs(title = "Salah et al., PCA by dose")
#> using ntop=500 top features by variance

pca_donors | pca_dose
```

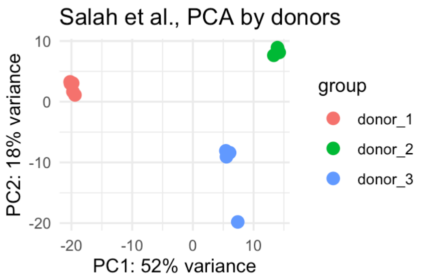


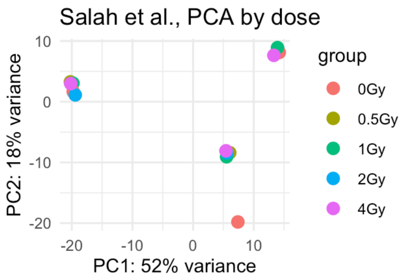


We quickly inspect a heatmap of the highly variable genes - as we have strong donor effect, we will account for it (removing that via `removeBatchEffect()`) for better visualization. The uncorrected raw counts will be used for the subsequent steps.

```
vsd_corrected <- vsd_salah_6h
assay(vsd_corrected) <- limma::removeBatchEffect(assay(vsd_corrected), batch = vsd_corrected$D
#> design matrix of interest not specified. Assuming a one-group experiment.
doses <- as.numeric(sub("Gy.*", "", colnames(vsd_corrected)))
col_order <- order(doses)
vsd_corrected <- vsd_corrected[, col_order]
colnames(vsd_corrected)
#>  [1] "0Gy-6hrs-do1"   "0Gy-6hrs-do2"   "0Gy-6hrs-do3"   "0.5Gy-6hrs-do1"
#>  [5] "0.5Gy-6hrs-do2" "0.5Gy-6hrs-do3" "1Gy-6hrs-do1"   "1Gy-6hrs-do2"
#>  [9] "1Gy-6hrs-do3"   "2Gy-6hrs-do1"   "2Gy-6hrs-do2"   "2Gy-6hrs-do3"
#> [13] "4Gy-6hrs-do1"   "4Gy-6hrs-do2"   "4Gy-6hrs-do3"

top_n <- 50
rv <- rowVars(assay(vsd_corrected))
top_genes <- order(rv, decreasing = TRUE)[1:top_n]
mat <- assay(vsd_corrected)[top_genes, ]
mat_scaled <- t(scale(t(mat)))
annotation <- as.data.frame(colData(vsd_corrected)[, c("Sex","Dose"), drop = FALSE])
Heatmap(mat_scaled,
        name = "expression",
        cluster_columns = FALSE,
        top_annotation = HeatmapAnnotation(df = annotation))
```

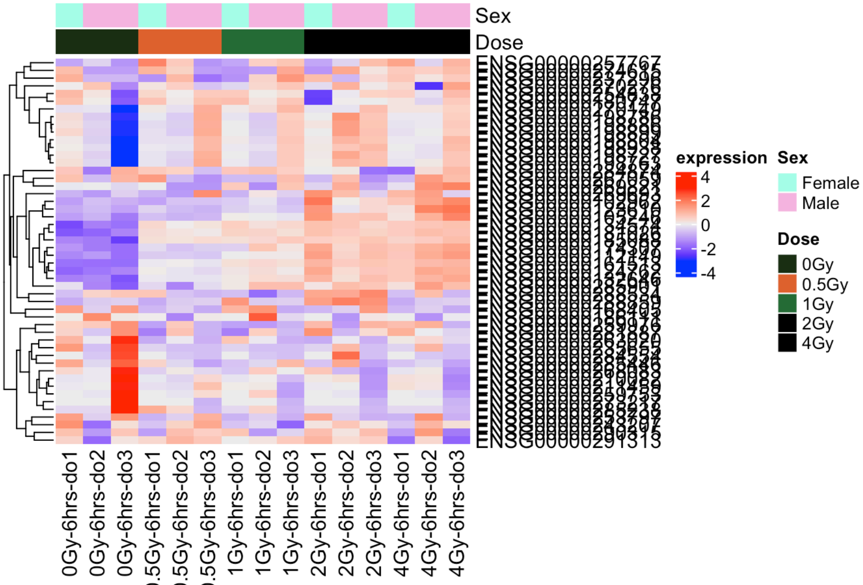


The columns of the heatmap are left ordered by dose.

Afterwards, we proceed with the modeling and testing step with DESeq2.

```r
dds_salah_6h
#> class: DESeqDataSet
#> dim: 62646 15
#> metadata(9): tximetaInfo quantInfo ... DoReMiTra version
#> assays(3): counts abundance avgTxLength
#> rownames(62646): ENSG00000000003 ENSG00000000005 ... ENSG00000293599
#>   ENSG00000293600
#> rowData names(3): gene_id tx_ids SYMBOL
#> colnames(15): 0.5Gy-6hrs-do1 1Gy-6hrs-do1 ... 2Gy-6hrs-do3 4Gy-6hrs-do3
#> colData names(10): Dose Time_point ... Exp_setting Platform

design(dds_salah_6h)<- ~Donors + Dose
dds_salah_6h <- DESeq(dds_salah_6h)
#> estimating size factors
#> using 'avgTxLength' from assays(dds), correcting for library size
#> estimating dispersions
#> gene-wise dispersion estimates
#> mean-dispersion relationship
#> final dispersion estimates
#> fitting model and testing
resultsNames(dds_salah_6h)
#> [1] "Intercept"                 "Donors_donor_2_vs_donor_1"
#> [3] "Donors_donor_3_vs_donor_1" "Dose_0.5Gy_vs_0Gy"
#> [5] "Dose_1Gy_vs_0Gy"           "Dose_2Gy_vs_0Gy"
#> [7] "Dose_4Gy_vs_0Gy"

# we will focus on one contrast here, 2Gy vs 0Gy
results_salah<- results(dds_salah_6h,
                        name = "Dose_2Gy_vs_0Gy",
                        alpha = 0.05)
summary(results_salah)
#>
#> out of 30201 with nonzero total read count
#> adjusted p-value < 0.05
#> LFC > 0 (up)       : 197, 0.65%
#> LFC < 0 (down)     : 77, 0.25%
#> outliers [1]       : 0, 0%
#> low counts [2]     : 14915, 49%
#> (mean count < 7)
#> [1] see 'cooksCutoff' argument of ?results
#> [2] see 'independentFiltering' argument of ?results
results_salah_shrink <- lfcShrink(dds_salah_6h,
                                  res = results_salah,
                                  coef = "Dose_2Gy_vs_0Gy")
#> using 'apeglm' for LFC shrinkage. If used in published research, please cite:
#>     Zhu, A., Ibrahim, J.G., Love, M.I. (2018) Heavy-tailed prior distributions for
#>     sequence count data: removing the noise and preserving large differences.
#>     Bioinformatics. https://doi.org/10.1093/bioinformatics/bty895

result_salah_sig <- as.data.frame(results_salah_shrink)
result_salah_sig <-
  result_salah_sig[!is.na(results_salah_shrink$padj) & result_salah_sig$padj <= 0.05, ]

# adding SYMBOL column
result_salah_sig$SYMBOL <- mapIds(
  org.Hs.eg.db,
  keys = rownames(result_salah_sig),
  column = "SYMBOL",
  keytype = "ENSEMBL",
  multiVals = "first"
)
#> 'select()' returned 1:many mapping between keys and columns

result_salah_sig |>
  head(10) |>
  knitr::kable()
```

| | baseMean | log2FoldChange | lfcSE | pvalue | padj | SYMBOL |
|---|---|---|---|---|---|---|
| ENSG00000009413 | 1466.8788 | 0.8194507 | 0.2331190 | 0.0000219 | 0.0032261 | REV3L |
| ENSG00000010818 | 3634.3748 | 0.6828260 | 0.1626106 | 0.0000016 | 0.0003454 | HIVEP2 |
| ENSG00000012779 | 2322.2113 | 0.3859968 | 0.1307390 | 0.0003284 | 0.0236759 | ALOX5 |
| ENSG00000013810 | 1421.9630 | -0.5529292 | 0.1858397 | 0.0001786 | 0.0156867 | TACC3 |
| ENSG00000023445 | 2664.3304 | 1.0234267 | 0.2760269 | 0.0000082 | 0.0014388 | BIRC3 |
| ENSG00000023892 | 1837.3742 | -0.5147620 | 0.1777516 | 0.0002644 | 0.0205193 | DEF6 |
| ENSG00000026103 | 802.8594 | 1.2610716 | 0.2076139 | 0.0000000 | 0.0000000 | FAS |
| ENSG00000033178 | 781.7486 | 0.8676891 | 0.2883771 | 0.0001036 | 0.0101479 | UBA6 |
| ENSG00000035664 | 499.3658 | -1.0199059 | 0.2264002 | 0.0000003 | 0.0000846 | DAPK2 |
| ENSG00000054148 | 223.0434 | 2.3754658 | 0.1837554 | 0.0000000 | 0.0000000 | PHPT1 |

We can also visualize the up and down regulated genes via a volcano plot

```
mosdef::de_volcano(
  res_de = results_salah_shrink,
  mapping = "org.Hs.eg.db",
  labeled_genes = 50,
  logfc_cutoff = 0
)
#> 'select()' returned 1:many mapping between keys and columns
#> Warning: Removed 32445 rows containing missing values or values outside the scale range
#> (`geom_point()`).
#> Warning: Removed 62597 rows containing missing values or values outside the scale range
#> (`geom_text_repel()`).
```

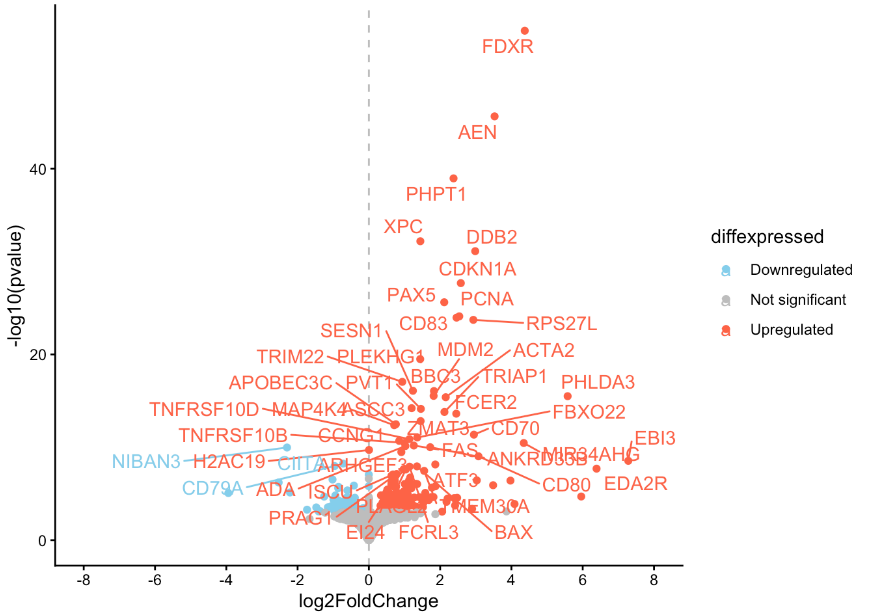


As we can see here, among the highest X-ray sensitive genes upon 2Gy exposure we can find FDXR, CDKN1A, AEN, and CD83.

From this point, we can also perform a functional enrichment analysis to see what pathways are regulated

```
res_enrich_salah <- enrichGO(result_salah_sig$SYMBOL,
                             keyType = "SYMBOL",
                             ont = "BP",
                             OrgDb = "org.Hs.eg.db",
                             pvalueCutoff = 0.05,
                             pAdjustMethod = "BH",
                             qvalueCutoff = 0.2,
                             minGSSize = 10,
                             maxGSSize = 500
)

res_enrich_salah_sig <- res_enrich_salah@result[res_enrich_salah@result$p.adjust <= 0.05, ]
res_enrich_salah_sig |>
  head(20) |>
  knitr::kable()
```

| | ID | Description | GeneRatio | BgRatio | RichFactor | FoldEnrichment | zScore | pvalue | p.adjus |
|---|---|---|---|---|---|---|---|---|---|
| GO:0072331 | GO:0072331 | signal transduction by p53 class mediator | 19/239 | 196/18860 | 0.0969388 | 7.649646 | 10.601833 | 0 | 0.0e+0 |
| GO:0097193 | GO:0097193 | intrinsic apoptotic signaling pathway | 24/239 | 335/18860 | 0.0716418 | 5.653407 | 9.735787 | 0 | 0.0e+0 |
| GO:0030098 | GO:0030098 | lymphocyte differentiation | 26/239 | 441/18860 | 0.0589569 | 4.652416 | 8.792721 | 0 | 1.0e-0 |
| GO:0072332 | GO:0072332 | intrinsic apoptotic signaling pathway by p53 class mediator | 13/239 | 88/18860 | 0.1477273 | 11.657474 | 11.352647 | 0 | 1.0e-0 |

| | ID | Description | GeneRatio | BgRatio | RichFactor | FoldEnrichment | zScore | pvalue | p.adjus |
|---|---|---|---|---|---|---|---|---|---|
| GO:0070661 | GO:0070661 | leukocyte proliferation | 23/239 | 356/18860 | 0.0646067 | 5.098256 | 8.843986 | 0 | 1.0e-0 |
| GO:0046651 | GO:0046651 | lymphocyte proliferation | 21/239 | 312/18860 | 0.0673077 | 5.311394 | 8.699679 | 0 | 3.0e-0 |
| GO:0071887 | GO:0071887 | leukocyte apoptotic process | 14/239 | 125/18860 | 0.1120000 | 8.838159 | 9.960910 | 0 | 3.0e-0 |
| GO:0032943 | GO:0032943 | mononuclear cell proliferation | 21/239 | 319/18860 | 0.0658307 | 5.194843 | 8.560541 | 0 | 3.0e-0 |
| GO:0008630 | GO:0008630 | intrinsic apoptotic signaling pathway in response to DNA damage | 13/239 | 111/18860 | 0.1171171 | 9.241962 | 9.866425 | 0 | 6.0e-0 |
| GO:0009411 | GO:0009411 | response to UV | 15/239 | 159/18860 | 0.0943396 | 7.444541 | 9.245170 | 0 | 6.0e-0 |
| GO:0070231 | GO:0070231 | T cell apoptotic process | 10/239 | 61/18860 | 0.1639344 | 12.936415 | 10.578601 | 0 | 1.4e-0 |
| GO:0071214 | GO:0071214 | cellular response to abiotic stimulus | 21/239 | 355/18860 | 0.0591549 | 4.668042 | 7.904250 | 0 | 1.4e-0 |
| GO:0104004 | GO:0104004 | cellular response to environmental stimulus | 21/239 | 355/18860 | 0.0591549 | 4.668042 | 7.904250 | 0 | 1.4e-0 |
| GO:0030330 | GO:0030330 | DNA damage response, signal transduction by p53 class mediator | 11/239 | 82/18860 | 0.1341463 | 10.585774 | 9.855218 | 0 | 1.6e-0 |
| GO:0046631 | GO:0046631 | alpha-beta T cell activation | 16/239 | 206/18860 | 0.0776699 | 6.129098 | 8.385817 | 0 | 1.9e-0 |
| GO:0070227 | GO:0070227 | lymphocyte apoptotic process | 11/239 | 85/18860 | 0.1294118 | 10.212158 | 9.643566 | 0 | 2.1e-0 |
| GO:0034644 | GO:0034644 | cellular response to UV | 11/239 | 92/18860 | 0.1195652 | 9.435146 | 9.188282 | 0 | 4.5e-0 |
| GO:0070663 | GO:0070663 | regulation of leukocyte proliferation | 18/239 | 284/18860 | 0.0633803 | 5.001473 | 7.697679 | 0 | 4.5e-0 |
| GO:0009314 | GO:0009314 | response to radiation | 23/239 | 462/18860 | 0.0497835 | 3.928526 | 7.220073 | 0 | 4.5e-0 |
| GO:0001776 | GO:0001776 | leukocyte homeostasis | 12/239 | 118/18860 | 0.1016949 | 8.024963 | 8.672291 | 0 | 5.7e-0 |

Now we will also visualize the top 10 regulated pathways.

```
# create GeneTonic object
anno_df <- mosdef::get_annotation_orgdb(
  de_container = dds_salah,
  orgdb_package = "org.Hs.eg.db",
  id_type = "ENSEMBL"
)
#> 'select()' returned 1:many mapping between keys and columns

gtl_salah_6h <- GeneTonic::GeneTonicList(
  dds = dds_salah_6h,
  res_de = results_salah,
  res_enrich = shake_enrichResult(res_enrich_salah),
  annotation_obj = anno_df
)
#> Found 3335 gene sets in `enrichResult` object, of which 276 are significant.
#> Converting for usage in GeneTonic...
#> ---------------------------------
#> ----- GeneTonicList object ------
#> ---------------------------------
#>
#> ----- dds object -----
#> Providing an expression object (as DESeqDataset) of 62646 features over 15 samples
```

```
#>
#> ----- res_de object -----
#> Providing a DE result object (as DESeqResults), 62646 features tested, 274 found as DE
#> Upregulated:     197
#> Downregulated:   77
#>
#> ----- res_enrich object -----
#> Providing an enrichment result object, 3335 reported
#>
#> ----- annotation_obj object -----
#> Providing an annotation object of 62646 features with information on 2 identifier types

# preparing the matrix for heatmap
scored_mat <- GeneTonic::gs_scores(se = vsd_corrected, gtl = gtl_salah_6h)

anno <- as.data.frame(colData(vsd_corrected)[, c("Sex", "Dose")])
dose_colors <- colorRampPalette(c("lightyellow", "gold", "orange", "red"))(5)
names(dose_colors)<- c("0Gy", "0.5Gy", "1Gy", "2Gy", "4Gy")
anno_colors <- list(
  Sex = c(Female = "pink", Male = "blue"),
  Dose = dose_colors
)
column_annotation <- HeatmapAnnotation(
  Sex = anno$Sex,
  Dose = anno$Dose,
  col = anno_colors, show_legend = TRUE
)

col_fun <- colorRamp2(
  c(-2, 0, 2),
  c("blue", "white", "red")
)

ht <- ComplexHeatmap::Heatmap(scored_mat[1:20,],
  name = "Expression",
  top_annotation = column_annotation,
  cluster_columns = FALSE, # Optional: Cluster columns
  cluster_rows = FALSE, # Optional: Cluster rows
  show_row_names = TRUE,
  show_column_names = TRUE,
  row_dend_side = "left",
  row_names_side = "left",
  row_names_gp = gpar(fontsize = 5, fontface= "bold" ),
  width = unit(7, "cm"),
  heatmap_height = unit(12, "cm"),
  column_names_gp = gpar(fontsize = 7),
  column_names_rot = 90,
  column_title = "Donors",
  column_title_side = "bottom",
  show_heatmap_legend = TRUE,
  col = col_fun
)

draw(ht, heatmap_legend_side = "right", annotation_legend_side = "right")
```

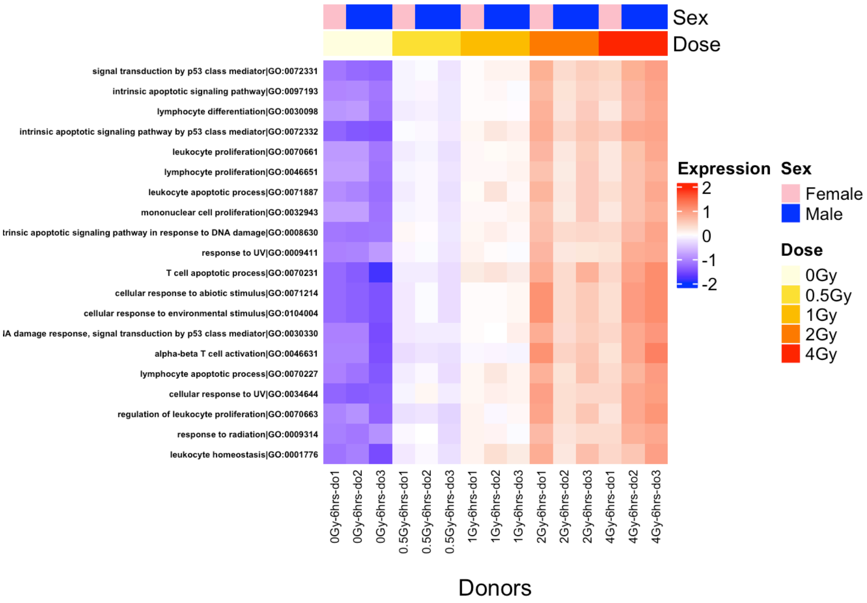


TODO; We can see here that among the top 20 regulated pathways we can find processes such as DNA damage repair, immune response, cellular response to UV, and some other associated GO terms.

#### 7.2 Working on the gamma ray dataset (Paul et al. 2013)

We might also want to compare Xray signature to gamma ray signatures, to see if both ionizing radiation types have shared and distinct transcriptional responses we will therefore focus on the `se_paul` dataset, since it has the same time point of `se_salah` (6 hours after irradiation).

In this `se_paul` dataset, Age and Sex should be accounted for to capture signals only in response to radiation

```
# for comparability, we will only focus on 6hr timepoint
se_paul_6h <- se_paul[, colData(se_paul)$Time_point == "6hr"]
# since it is a repeated measurements experiment, we will create a pseudo-donor factor to acc

expr_values <- assay(se_paul_6h)
# remove probes with NA values
expr_values <- expr_values[rowSums(is.na(expr_values)) == 0, ]

# remove probes with zero variance
expr_values <- expr_values[apply(expr_values, 1, sd) != 0, ]

# extracting metadata
meta_df <- as.data.frame(colData(se_paul_6h))

# number of donors
n_donors <- 5
# number of doses per donor
n_doses <- 5

# create pseudo-donor factor
meta_df$Donors <- factor(rep(1:n_donors, each = n_doses))

# check
table(meta_df$Donors)
#> 
#> 1 2 3 4 5 
#> 5 5 5 5 5 

# handle the covariates as factors
meta_df$Dose <- factor(meta_df$Dose)
meta_df$Dose <- relevel(meta_df$Dose, ref = "0Gy")
meta_df$Sex  <- factor(meta_df$Sex)
meta_df$Age <- as.numeric(gsub("[^0-9]", "", meta_df$Age))

# building the design matrix
design_paul <- model.matrix(~ Age + Sex + Dose, data = meta_df)
design_paul
#>            (Intercept) Age Sexmale Dose0.5Gy Dose2Gy Dose5Gy Dose8Gy
#> GSM1080589           1  39       1         1       0       0       0
#> GSM1080590           1  39       1         0       0       0       0
#> GSM1080591           1  39       1         0       1       0       0
#> GSM1080592           1  39       1         0       0       1       0
#> GSM1080593           1  39       1         0       0       0       1
#> GSM1080594           1  20       0         1       0       0       0
#> GSM1080595           1  20       0         0       0       0       0
#> GSM1080596           1  20       0         0       1       0       0
#> GSM1080597           1  20       0         0       0       1       0
#> GSM1080598           1  20       0         0       0       0       1
#> GSM1080599           1  53       1         1       0       0       0
#> GSM1080600           1  53       1         0       0       0       0
#> GSM1080601           1  53       1         0       1       0       0
#> GSM1080602           1  53       1         0       0       1       0
#> GSM1080603           1  53       1         0       0       0       1
#> GSM1080604           1  35       0         1       0       0       0
#> GSM1080605           1  35       0         0       0       0       0
#> GSM1080606           1  35       0         0       1       0       0
#> GSM1080607           1  35       0         0       0       1       0
#> GSM1080608           1  35       0         0       0       0       1
#> GSM1080609           1  36       0         1       0       0       0
#> GSM1080610           1  36       0         0       0       0       0
#> GSM1080611           1  36       0         0       1       0       0
#> GSM1080612           1  36       0         0       0       1       0
#> GSM1080613           1  36       0         0       0       0       1
#> attr(,"assign")
#> [1] 0 1 2 3 3 3 3
#> attr(,"contrasts")
#> attr(,"contrasts")$Sex
#> [1] "contr.treatment"
#> 
#> attr(,"contrasts")$Dose
#> [1] "contr.treatment"

# blocking for the donor effect
corfit_paul <- duplicateCorrelation(expr_values, design_paul, block = meta_df$Donors)
```

```
# fitting the model
fit_paul <- lmFit(expr_values, design_paul, block = meta_df$Patient, correlation = corfit_paul

contrast.matrix_paul <- makeContrasts(Dose2Gy, levels = design_paul)
#> Renaming (Intercept) to Intercept
fit2_paul <- contrasts.fit(fit_paul, contrast.matrix_paul)
fit2_paul <- eBayes(fit2_paul)

# extracting the results
results_paul <- topTable(fit2_paul, number = Inf)
results_paul_sig <- results_paul[ results_paul$adj.P.Val<= 0.05,]

results_paul_sig |>
  head(10) |>
  knitr::kable()
```

| | logFC | AveExpr | t | P.Value | adj.P.Val | B |
|---|---|---|---|---|---|---|
| FDXR | 5.114015 | 11.828418 | 45.28673 | 0 | 0 | 31.89617 |
| DDB2 | 2.971409 | 11.710745 | 28.29392 | 0 | 0 | 27.42778 |
| CD70 | 3.501244 | 9.363679 | 23.68030 | 0 | 0 | 25.17089 |
| PCNA | 2.691635 | 9.432219 | 21.75109 | 0 | 0 | 23.99659 |
| _A_24_P127462 | 2.900812 | 8.107424 | 21.66597 | 0 | 0 | 23.94103 |
| RPS27L | 1.994590 | 11.034623 | 18.90364 | 0 | 0 | 21.94062 |
| GADD45A | 2.100923 | 9.198538 | 17.26089 | 0 | 0 | 20.54319 |
| _A_32_P85230 | 3.472482 | 8.768788 | 17.00659 | 0 | 0 | 20.31096 |
| SESN1_A_23_P93562 | 1.806004 | 8.151917 | 16.02157 | 0 | 0 | 19.36746 |
| DCP1B | 1.271772 | 9.040659 | 15.50149 | 0 | 0 | 18.83951 |

Here, we will extract the gene symbol part without the probe id so we can compare the genes with the Salah_2025 dataset

```
results_paul_sig$SYMBOL <- sub("_.*", "", rownames(results_paul_sig))

# again, we can also check this effect at the pathway level
res_enrich_paul <- enrichGO(results_paul_sig$SYMBOL,
                            keyType = "SYMBOL",
                            ont = "BP",
                            OrgDb = "org.Hs.eg.db",
                            pvalueCutoff = 0.05,
                            pAdjustMethod = "BH",
                            qvalueCutoff = 0.2,
                            minGSSize = 10,
                            maxGSSize = 500
)

res_enrich_paul_sig <- res_enrich_paul@result[res_enrich_paul@result$p.adjust <= 0.05, ]

res_enrich_paul_sig |>
  head(20) |>
  knitr::kable()
```

| | ID | Description | GeneRatio | BgRatio | RichFactor | FoldEnrichment | zScore | pvalue | p.adju |
|---|---|---|---|---|---|---|---|---|---|
| GO:0097193 | GO:0097193 | intrinsic apoptotic signaling pathway | 18/95 | 335/18860 | 0.0537313 | 10.667086 | 12.702398 | 0 | 0e+0 |
| GO:0071214 | GO:0071214 | cellular response to abiotic stimulus | 18/95 | 355/18860 | 0.0507042 | 10.066123 | 12.269817 | 0 | 0e+0 |
| GO:0104004 | GO:0104004 | cellular response to environmental stimulus | 18/95 | 355/18860 | 0.0507042 | 10.066123 | 12.269817 | 0 | 0e+0 |
| GO:0034644 | GO:0034644 | cellular response to UV | 11/95 | 92/18860 | 0.1195652 | 23.736842 | 15.554710 | 0 | 0e+0 |
| GO:0009411 | GO:0009411 | response to UV | 13/95 | 159/18860 | 0.0817610 | 16.231711 | 13.723399 | 0 | 0e+0 |
| GO:0071478 | GO:0071478 | cellular response to radiation | 13/95 | 188/18860 | 0.0691489 | 13.727884 | 12.479194 | 0 | 0e+0 |

| | ID | Description | GeneRatio | BgRatio | RichFactor | FoldEnrichment | zScore | pvalue | p.adju |
|---|---|---|---|---|---|---|---|---|---|
| GO:0009314 | GO:0009314 | response to radiation | 18/95 | 462/18860 | 0.0389610 | 7.734792 | 10.428136 | 0 | 0e+0 |
| GO:2001242 | GO:2001242 | regulation of intrinsic apoptotic signaling pathway | 13/95 | 195/18860 | 0.0666667 | 13.235088 | 12.219607 | 0 | 0e+0 |
| GO:0072331 | GO:0072331 | signal transduction by p53 class mediator | 13/95 | 196/18860 | 0.0663265 | 13.167562 | 12.183613 | 0 | 0e+0 |
| GO:0072332 | GO:0072332 | intrinsic apoptotic signaling pathway by p53 class mediator | 10/95 | 88/18860 | 0.1136364 | 22.559809 | 14.423738 | 0 | 0e+0 |
| GO:0071482 | GO:0071482 | cellular response to light stimulus | 11/95 | 124/18860 | 0.0887097 | 17.611205 | 13.204453 | 0 | 0e+0 |
| GO:0008630 | GO:0008630 | intrinsic apoptotic signaling pathway in response to DNA damage | 10/95 | 111/18860 | 0.0900901 | 17.885254 | 12.694830 | 0 | 0e+0 |
| GO:0030330 | GO:0030330 | DNA damage response, signal transduction by p53 class mediator | 9/95 | 82/18860 | 0.1097561 | 21.789474 | 13.423711 | 0 | 1e-0 |
| GO:0070227 | GO:0070227 | lymphocyte apoptotic process | 9/95 | 85/18860 | 0.1058824 | 21.020433 | 13.162543 | 0 | 1e-0 |
| GO:0071887 | GO:0071887 | leukocyte apoptotic process | 10/95 | 125/18860 | 0.0800000 | 15.882105 | 11.877892 | 0 | 1e-0 |
| GO:0070231 | GO:0070231 | T cell apoptotic process | 8/95 | 61/18860 | 0.1311475 | 26.036238 | 13.935208 | 0 | 1e-0 |
| GO:0070228 | GO:0070228 | regulation of lymphocyte apoptotic process | 8/95 | 63/18860 | 0.1269841 | 25.209691 | 13.695002 | 0 | 1e-0 |
| GO:2000106 | GO:2000106 | regulation of leukocyte apoptotic process | 9/95 | 97/18860 | 0.0927835 | 18.419967 | 12.238520 | 0 | 2e-0 |
| GO:0009416 | GO:0009416 | response to light stimulus | 14/95 | 341/18860 | 0.0410557 | 8.150640 | 9.481133 | 0 | 2e-0 |
| GO:2001233 | GO:2001233 | regulation of apoptotic signaling pathway | 15/95 | 406/18860 | 0.0369458 | 7.334716 | 9.181038 | 0 | 2e-0 |

Similarly to X-ray dataset from `se_salah`, many of the regulated pathways were involved in processes such as DNA damage repair, immune response, and response to UV.

### 8 Integrating the results from both studies

Checking the shared genes between both radiation types, at the same timepoint and radiation dose

```
result_salah_sig<- na.omit(result_salah_sig)

# creating a simple venn diagram first
venn.plot <- venn.diagram(
  x = list(
    Salah = result_salah_sig$SYMBOL,
    Paul = results_paul_sig$SYMBOL
  ),
  fill = c("#8da0cb", "orange"),
  alpha = 0.6,
```

```
  cat.cex = 1.2,
  cex = 1.5,
  main = "Overlap of DE genes - Shared and distinct signatures",
  filename = NULL,
  disable.logging = TRUE)
#> INFO [2025-12-12 16:46:10] $x
#> INFO [2025-12-12 16:46:10] list(Salah = result_salah_sig$SYMBOL, Paul = results_paul_sig$SY
#> INFO [2025-12-12 16:46:10]
#> INFO [2025-12-12 16:46:10] $fill
#> INFO [2025-12-12 16:46:10] c("#8da0cb", "orange")
#> INFO [2025-12-12 16:46:10]
#> INFO [2025-12-12 16:46:10] $alpha
#> INFO [2025-12-12 16:46:10] [1] 0.6
#> INFO [2025-12-12 16:46:10]
#> INFO [2025-12-12 16:46:10] $cat.cex
#> INFO [2025-12-12 16:46:10] [1] 1.2
#> INFO [2025-12-12 16:46:10]
#> INFO [2025-12-12 16:46:10] $cex
#> INFO [2025-12-12 16:46:10] [1] 1.5
#> INFO [2025-12-12 16:46:10]
#> INFO [2025-12-12 16:46:10] $main
#> INFO [2025-12-12 16:46:10] [1] "Overlap of DE genes - Shared and distinct signatures"
#> INFO [2025-12-12 16:46:10]
#> INFO [2025-12-12 16:46:10] $filename
#> INFO [2025-12-12 16:46:10] NULL
#> INFO [2025-12-12 16:46:10]
#> INFO [2025-12-12 16:46:10] $disable.logging
#> INFO [2025-12-12 16:46:10] [1] TRUE
#> INFO [2025-12-12 16:46:10]

grid::grid.draw(venn.plot)
```

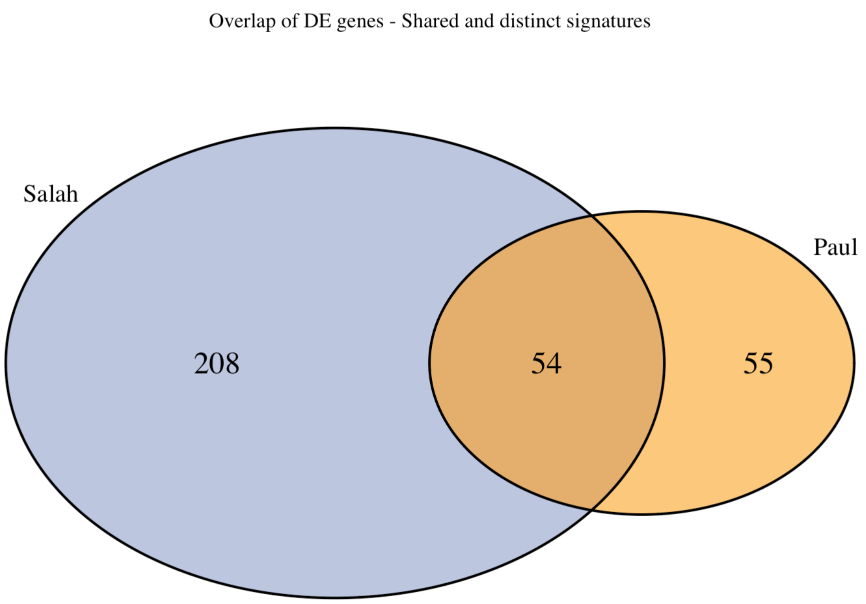


```
# shared signature between X and gamma rays
intersect(result_salah_sig$SYMBOL, results_paul_sig$SYMBOL)
#>  [1] "REV3L"     "FAS"       "PHPT1"     "BCL3"      "NFKB2"     "SESN1"
#>  [7] "BAX"       "IZUMO4"    "CD40"      "RELB"      "EBI3"      "BBC3"
#> [13] "CD79A"     "ACTA2"     "SYNGR2"    "CD83"      "PTP4A1"    "ASCC3"
#> [19] "TMEM30A"   "CCNG1"     "GADD45A"   "TNFRSF10B" "CDKN1A"    "CD70"
#> [25] "PLAGL2"    "CCR7"      "GDF15"     "TRIM22"    "PCNA"      "DDB2"
#> [31] "MDM2"      "DRAM1"     "MYC"       "PMAIP1"    "FGD2"      "EI24"
#> [37] "DCP1B"     "BCL2L11"   "XPC"       "FDXR"      "IER5"      "ARHGEF3"
#> [43] "FBXO22"    "VWCE"      "POLH"      "PPM1D"     "TRIAP1"    "BCL2"
#> [49] "ZMAT3"     "RPS27L"    "ADA"       "ANXA4"     "APOBEC3C"  "MARCKS"

# gamma-ray only signature
setdiff(results_paul_sig$SYMBOL, result_salah_sig$SYMBOL)
#>  [1] ""          "FAM129C"   "ANKRA2"    "PLK3"      "C12orf5"   "CCDC90B"
#>  [7] "MGAT3"     "LIG1"      "C17orf89"  "ZNF337"    "METTL7A"   "PLK2"
#> [13] "IL21R"     "SERPINB1"  "TTC21A"    "GPR84"     "TNIP1"     "LY9"
#> [19] "C3orf26"   "SLC7A6"    "APOBEC3F"  "THOP1"     "RGS12"     "SCRIB"
#> [25] "PTGS2"     "GHRL"      "PYCR1"     "TNFAIP6"   "P2RX5"     "ZMIZ2"
#> [31] "PLEKHF1"   "SAC3D1"    "FAM136A"   "SGK223"    "UPB1"      "METTL12"
#> [37] "RFTN1"     "IL8"       "TNFRSF12A" "ID2"       "BTG3"      "JAK3"
#> [43] "MR1"       "ZNF195"    "MRPL49"    "TMEM205"   "TM7SF3"    "TESK2"
#> [49] "SNHG12"    "FBXW7"     "NAP1L1"    "TP53TG1"   "POPDC2"    "TSPAN32"
```

```
#> [55] "ERN1"

# X-ray only signature
setdiff(result_salah_sig$SYMBOL, results_paul_sig$SYMBOL)
#>   [1] "HIVEP2"       "ALOX5"        "TACC3"        "BIRC3"        "DEF6"
#>   [6] "UBA6"         "DAPK2"        "PRDM1"        "RASGRF1"      "TBXAS1"
#>  [11] "CALCRL"       "IDI1"         "MAP4K4"       "ACTN1"        "ACAP1"
#>  [16] "ZNF37A"       "PLD1"         "TRAF4"        "TIGAR"        "TP53INP2"
#>  [21] "EPB41L2"      "PCNP"         "COBLL1"       "PIBF1"        "CYLD"
#>  [26] "ATRN"         "SLC8B1"       "HIVEP1"       "PPIL2"        "APOBEC3H"
#>  [31] "CSF2RB"       "HDAC10"       "HIF1A"        "PPP4R3A"      "CCM2L"
#>  [36] "TM9SF4"       "RBBP8"        "RENBP"        "JADE3"        "ACP5"
#>  [41] "N4BP1"        "CCL22"        "GGA2"         "NBN"          "FCER2"
#>  [46] "ASF1B"        "DBP"          "GSK3A"        "SNX8"         "TNFSF8"
#>  [51] "LIPA"         "GBF1"         "UNC119"       "NFKB1"        "DBN1"
#>  [56] "GORASP1"      "IFIH1"        "RALGPS2"      "RNF19B"       "WLS"
#>  [61] "SIPA1L2"      "TNFSF4"       "PLAGL1"       "TNFAIP3"      "SGK1"
#>  [66] "ATL2"         "PANK3"        "PLEKHG1"      "CD80"         "ZC3H7A"
#>  [71] "MRM2"         "KCNJ2"        "PI3"          "PLCG1"        "MIF4GD"
#>  [76] "TNFSF9"       "MED1"         "CD93"         "IQCN"         "PNPLA7"
#>  [81] "HIP1R"        "EDA2R"        "TRAF3"        "NPHP4"        "PTGFRN"
#>  [86] "NOTCH2"       "IL2RA"        "GLS2"         "NHSL1"        "ISCU"
#>  [91] "LIMD2"        "IRF4"         "MMP7"         "ACVRL1"       "NKD1"
#>  [96] "FN3KRP"       "IGFBP4"       "SLC2A5"       "RGL1"         "GOLGA4"
#> [101] "LPP"          "TIFA"         "FEZ1"         "MERTK"        "NDUFAF6"
#> [106] "ZDHHC5"       "ZFAND3"       "MS4A1"        "ODR4"         "SNX22"
#> [111] "PANK4"        "RNF207"       "TSPAN33"      "B4GALT5"      "CD1D"
#> [116] "CD1C"         "CCAR2"        "SON"          "TNFRSF13C"    "PTGIR"
#> [121] "JAML"         "PCSK7"        "FCRL3"        "PWWP3A"       "ABCF3"
#> [126] "LRP5"         "MEGF6"        "PEA15"        "ATF3"         "REL"
#> [131] "SMC6"         "CDC42EP3"     "PYHIN1"       "UVSSA"        "SLBP"
#> [136] "ANKRD33B"     "CEP295"       "ABTB2"        "TCP11L2"      "NOLC1"
#> [141] "ZCCHC18"      "DPEP2"        "NIBAN3"       "TAPT1"        "C11orf24"
#> [146] "LRG1"         "WIPF2"        "PTGER4"       "PIK3CD"       "GPR82"
#> [151] "SYNPO"        "FRMD3"        "CEBPB"        "TRIB1"        "TNFRSF10D"
#> [156] "EIF1AX"       "PHLDA3"       "CKAP5"        "P2RY2"        "MLXIP"
#> [161] "GNG7"         "ACER2"        "PTPN11"       "CIITA"        "LACC1"
#> [166] "AEN"          "ZNF746"       "FRAT2"        "TSEN54"       "CCDC125"
#> [171] "TNFAIP2"      "L3MBTL1"      "IER5L"        "FAM111B"      "PAX5"
#> [176] "ZNF79"        "ZNF431"       "KIAA1671"     "SDAD1"        "BAZ1A"
#> [181] "MDM4"         "RUSC2"        "BRD2"         "IGKC"         "CEBPD"
#> [186] "LOC102723566" "FAM30A"       "MIR34AHG"     "TRAF3IP2-AS1" "MIR155HG"
#> [191] "TMEM250"      "IGKV1-33"     "NAIP"         "PVT1"         "IGKV1D-39"
#> [196] "CORO7"        "CCL4"         "PRAG1"        "CCL4L2"       "GTSE1-DT"
#> [201] "CCL3"         "ZNF229"       "LOC124901671" "SMIM10L3"     "H2AC19"
#> [206] "LOC100294145" "NSUN5P1"      "GTF2IP4"
```

Another way to display the shared and distinct regulation happening at the overall transcriptome level would be a logFC-logFC plot - of course, with the caveat of these two datasets originating from two different technology platforms (RNA-seq for `se_salah` and microarray for `se_paul`)

```
dim(results_salah_shrink)
#> [1] 62646     5
dim(results_paul)
#> [1] 14936     6

results_salah_shrink$SYMBOL <- mapIds(
  org.Hs.eg.db,
  keys = rownames(results_salah_shrink),
  column = "SYMBOL",
  keytype = "ENSEMBL",
  multiVals = "first"
)
#> 'select()' returned 1:many mapping between keys and columns

results_paul$SYMBOL <- rownames(results_paul)

merged_results <- merge.data.frame(x = as.data.frame(results_salah_shrink),
                                   y = results_paul,
                                   by = "SYMBOL")
merged_results$DE_salah <- merged_results$padj <= 0.05 & !is.na(merged_results$padj)
merged_results$DE_paul <- merged_results$adj.P.Val <= 0.05

ggplot(merged_results,
       aes(x = log2FoldChange, y = logFC, col = DE_salah)) +
  geom_point() +
  scale_color_manual(values = c(scales::alpha("black", 0.3), "firebrick")) +
  theme_bw() +
  labs(x = "logFC - RNA-seq",
       y = "logFC - microarray",
       title = "Highlighting DE genes from Salah et al.")
#> Warning: Removed 123 rows containing missing values or values outside the scale range
#> (`geom_point()`).
```

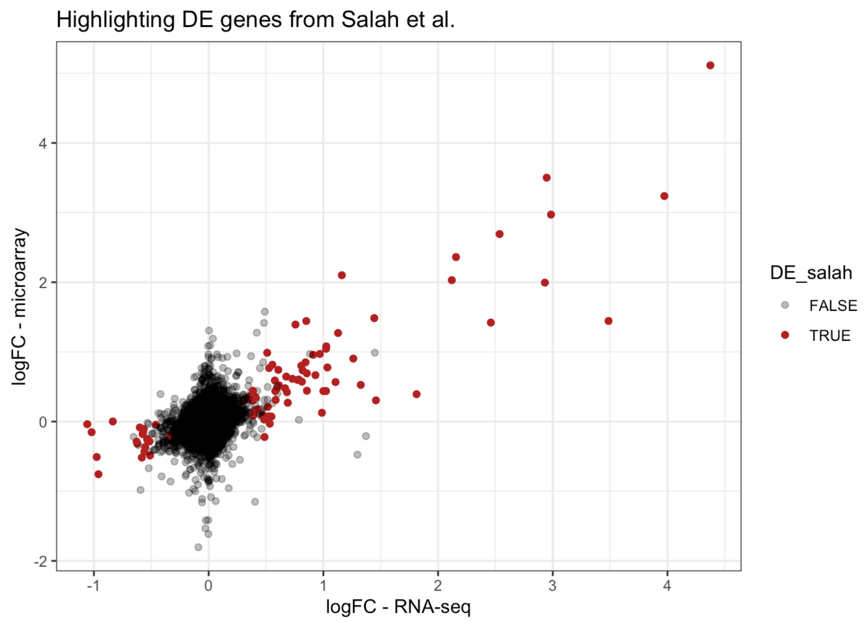


```r
ggplot(merged_results,
         aes(x = log2FoldChange, y = logFC, col = DE_paul)) +
  geom_point() +
  scale_color_manual(values = c(scales::alpha("black", 0.3), "firebrick")) +
  theme_bw() +
  labs(x = "logFC - RNA-seq",
       y = "logFC - microarray",
       title = "Highlighting DE genes from Paul et al.")
#> Warning: Removed 123 rows containing missing values or values outside the scale range
#> (`geom_point()`).
```

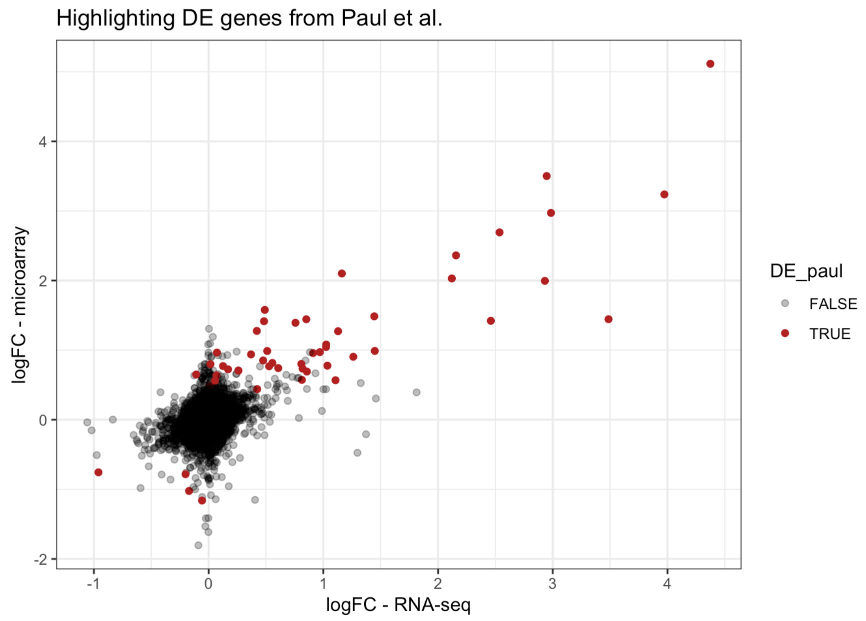


We can appreciate better how the gene expression regulation is to be assessed in quantitative nuances that overcome the dichotomization between significance or not, and the scatter plot of effect sizes can also better highlight the concordance in regulation direction.

One can also easily plot a geneset signature derived from one dataset (e.g. GO:0009411, response to UV, found in the microarray analysis), using the expression values of the other (Salah et al.).

```r
genes_uvresponse_paul <- res_enrich_paul_sig[res_enrich_paul_sig$ID == "GO:0009411", "geneID"]
  strsplit(split = "/") |>
  unlist()

genes_uvresponse_paul_ensembl <- mapIds(org.Hs.eg.db,
                                        keys = genes_uvresponse_paul,
                                        column = "ENSEMBL",
                                        keytype = "SYMBOL")
#> 'select()' returned 1:1 mapping between keys and columns

GeneTonic::gs_heatmap(se = vsd_corrected,
```

```
                      gtl = gtl_salah_6h,
                      genelist = genes_uvresponse_paul_ensembl,
                      plot_title = "Using the DE genes from Paul et al.")
```

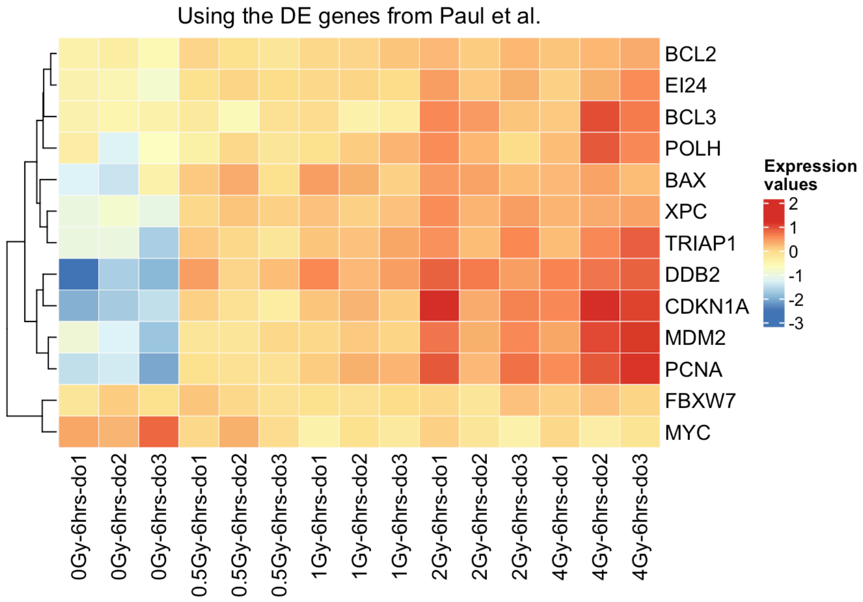


```
GeneTonic::gs_heatmap(se = vsd_corrected,
                      gtl = gtl_salah_6h,
                      geneset_id = "GO:0009411",
                      plot_title = "Using the original DE genes for UV response from Salah et
```

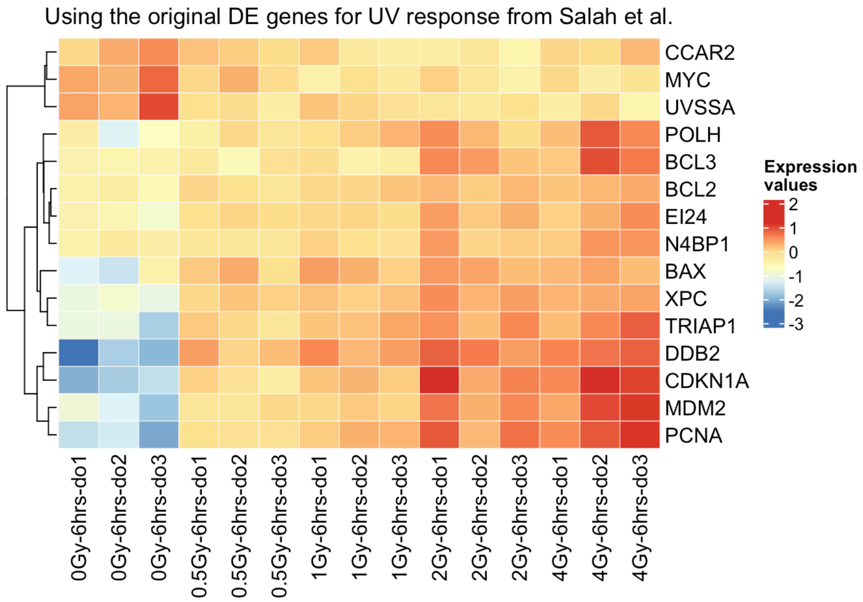


While most of the genes are shared, the higher power in DE detection from Salah et al. made it possible to identify a more comprehensive signature linked to this biological process.

Similarly to what we have just showcased, it is possible to compare other genesets and signatures across the individual datasets, e.g. apoptosis, response to DNA damage, and immune responses.

### 9 Wrapping up

This notebook can be further expanded to include additional datasets - e.g., scenarios where some discordant regulation is expected. Having `DoReMiTra` providing a number of them in this analysis-ready format makes it easy to explore them in detail, ultimately better understanding the transcriptional changes occurring upon radiation.

Overall, data integration and comparison through well-curated radiation transcriptomic datasets available through the `DoReMiTra` collection is useful in biomarker discovery upon radiation exposure.

#### Session info

```
sessionInfo()
#> R Under development (unstable) (2025-10-21 r88958)
#> Platform: x86_64-apple-darwin20
#> Running under: macOS Sequoia 15.6
#> 
#> Matrix products: default
#> BLAS:   /Library/Frameworks/R.framework/Versions/4.6-x86_64/Resources/lib/libRblas.0.dylib
#> LAPACK: /Library/Frameworks/R.framework/Versions/4.6-x86_64/Resources/lib/libRlapack.dylib;
#> 
#> locale:
#> [1] en_US.UTF-8/en_US.UTF-8/en_US.UTF-8/C/en_US.UTF-8/en_US.UTF-8
#> 
#> time zone: Europe/Berlin
#> tzcode source: internal
#> 
#> attached base packages:
#> [1] grid      stats4    stats     graphics  grDevices utils     datasets 
#> [8] methods   base     
#> 
#> other attached packages:
#>  [1] DoReMiTraExplorer_1.2.0     patchwork_1.3.2            
#>  [3] clusterProfiler_4.19.1      GeneTonic_3.5.0            
#>  [5] circlize_0.4.17             limma_3.67.0               
#>  [7] VennDiagram_1.7.3           futile.logger_1.4.3        
#>  [9] ComplexHeatmap_2.27.0       ggplot2_4.0.1              
#> [11] DESeq2_1.51.6               SummarizedExperiment_1.41.0
#> [13] MatrixGenerics_1.23.0       matrixStats_1.5.0          
#> [15] GenomicRanges_1.63.1        Seqinfo_1.1.0              
#> [17] org.Hs.eg.db_3.22.0         AnnotationDbi_1.73.0       
#> [19] IRanges_2.45.0              S4Vectors_0.49.0           
#> [21] Biobase_2.71.0              BiocGenerics_0.57.0        
#> [23] generics_0.1.4              mosdef_1.7.0               
#> [25] DoReMiTra_1.1.0            
#> 
#> loaded via a namespace (and not attached):
#>   [1] fs_1.6.6                    bitops_1.0-9               
#>   [3] enrichplot_1.31.0           httr_1.4.7                 
#>   [5] RColorBrewer_1.1-3          doParallel_1.0.17          
#>   [7] numDeriv_2016.8-1.1         dynamicTreeCut_1.63-1      
#>   [9] tippy_0.1.0                 tools_4.6.0                
#>  [11] R6_2.6.1                    DT_0.34.0                  
#>  [13] lazyeval_0.2.2              mgcv_1.9-4                 
#>  [15] apeglm_1.33.0               GetoptLong_1.1.0           
#>  [17] withr_3.0.2                 prettyunits_1.2.0          
#>  [19] gridExtra_2.3               textshaping_1.0.4          
#>  [21] cli_3.6.5                   formatR_1.14               
#>  [23] Cairo_1.7-0                 labeling_0.4.3             
#>  [25] sass_0.4.10                 topGO_2.63.0               
#>  [27] bs4Dash_2.3.5               mvtnorm_1.3-3              
#>  [29] S7_0.2.1                    ggridges_0.5.7             
#>  [31] goseq_1.63.0                Rsamtools_2.27.0           
#>  [33] systemfonts_1.3.1           yulab.utils_0.2.1          
#>  [35] gson_0.1.0                  txdbmaker_1.7.1            
#>  [37] svglite_2.2.2               DOSE_4.5.0                 
#>  [39] R.utils_2.13.0              scater_1.39.0              
#>  [41] dichromat_2.0-0.1           bbmle_1.0.25.1             
#>  [43] rstudioapi_0.17.1           RSQLite_2.4.5              
#>  [45] visNetwork_2.1.4            gridGraphics_0.5-1         
#>  [47] shape_1.4.6.1               BiocIO_1.21.0              
#>  [49] dplyr_1.1.4                 dendextend_1.19.1          
#>  [51] GO.db_3.22.0                Matrix_1.7-4               
#>  [53] ggbeeswarm_0.7.3            abind_1.4-8                
#>  [55] R.methodsS3_1.8.2           lifecycle_1.0.4            
#>  [57] yaml_2.3.12                 qvalue_2.43.0              
#>  [59] SparseArray_1.11.8          BiocFileCache_3.1.0        
#>  [61] blob_1.2.4                  promises_1.5.0             
#>  [63] bdsmatrix_1.3-7             ExperimentHub_3.1.0        
#>  [65] crayon_1.5.3                miniUI_0.1.2               
#>  [67] ggtangle_0.0.8              lattice_0.22-7             
#>  [69] beachmat_2.27.0             ComplexUpset_1.3.6         
#>  [71] cowplot_1.2.0               GenomicFeatures_1.63.1     
#>  [73] cigarillo_1.1.0             KEGGREST_1.51.1            
#>  [75] magick_2.9.0                pillar_1.11.1              
#>  [77] knitr_1.50                  fgsea_1.37.0               
#>  [79] rjson_0.2.23                codetools_0.2-20           
#>  [81] fastmatch_1.1-6             glue_1.8.0                 
#>  [83] ggiraph_0.9.2               ggfun_0.2.0                
#>  [85] fontLiberation_0.1.0        data.table_1.17.8          
#>  [87] vctrs_0.6.5                 png_0.1-8                  
```

```
#>  [89] treeio_1.35.0               gtable_0.3.6
#>  [91] emdbook_1.3.14              cachem_1.1.0
#>  [93] xfun_0.54                   S4Arrays_1.11.1
#>  [95] mime_0.13                   coda_0.19-4.1
#>  [97] SingleCellExperiment_1.33.0 iterators_1.0.14
#>  [99] statmod_1.5.1               nlme_3.1-168
#> [101] ggtree_4.1.1                bit64_4.6.0-1
#> [103] fontquiver_0.2.1            progress_1.2.3
#> [105] filelock_1.0.3              GenomeInfoDb_1.47.0
#> [107] bslib_0.9.0                 irlba_2.3.5.1
#> [109] vipor_0.4.7                 otel_0.2.0
#> [111] colorspace_2.1-2            DBI_1.2.3
#> [113] tidyselect_1.2.1            bit_4.6.0
#> [115] compiler_4.6.0              curl_7.0.0
#> [117] httr2_1.2.2                 graph_1.89.0
#> [119] BiocNeighbors_2.5.0         BiasedUrn_2.0.12
#> [121] SparseM_1.84-2              expm_1.0-0
#> [123] xml2_1.5.0                  plotly_4.11.0
#> [125] fontBitstreamVera_0.1.1     DelayedArray_0.37.0
#> [127] colourpicker_1.3.0          rtracklayer_1.71.0
#> [129] scales_1.4.0                rappdirs_0.3.3
#> [131] stringr_1.6.0               digest_0.6.39
#> [133] rmarkdown_2.30              XVector_0.51.0
#> [135] htmltools_0.5.9             pkgconfig_2.0.3
#> [137] dbplyr_2.5.1                fastmap_1.2.0
#> [139] rlang_1.1.6                 GlobalOptions_0.1.3
#> [141] htmlwidgets_1.6.4           UCSC.utils_1.7.0
#> [143] shiny_1.12.1                jquerylib_0.1.4
#> [145] farver_2.1.2                jsonlite_2.0.0
#> [147] BiocParallel_1.45.0         GOSemSim_2.37.0
#> [149] R.oo_1.27.1                 BiocSingular_1.27.1
#> [151] RCurl_1.98-1.17             magrittr_2.0.4
#> [153] kableExtra_1.4.0            scuttle_1.21.0
#> [155] ggplotify_0.1.3             Rcpp_1.1.0.8.1
#> [157] shinycssloaders_1.1.0       ape_5.8-1
#> [159] viridis_0.6.5               gdtools_0.4.4
#> [161] rintrojs_0.3.4              stringi_1.8.7
#> [163] MASS_7.3-65                 AnnotationHub_4.1.0
#> [165] plyr_1.8.9                  parallel_4.6.0
#> [167] ggrepel_0.9.6               Biostrings_2.79.2
#> [169] splines_4.6.0               hms_1.1.4
#> [171] geneLenDataBase_1.47.0      locfit_1.5-9.12
#> [173] igraph_2.2.1                ScaledMatrix_1.19.0
#> [175] reshape2_1.4.5              biomaRt_2.67.0
#> [177] futile.options_1.0.1        BiocVersion_3.23.1
#> [179] XML_3.99-0.20               evaluate_1.0.5
#> [181] lambda.r_1.2.4              BiocManager_1.30.27
#> [183] foreach_1.5.2               tweenr_2.0.3
#> [185] httpuv_1.6.16               backbone_3.0.2
#> [187] tidyr_1.3.1                 purrr_1.2.0
#> [189] polyclip_1.10-7             clue_0.3-66
#> [191] ggforce_0.5.0               rsvd_1.0.5
#> [193] xtable_1.8-4                restfulr_0.0.16
#> [195] tidytree_0.4.6              later_1.4.4
#> [197] viridisLite_0.4.2           tibble_3.3.0
#> [199] aplot_0.2.9                 beeswarm_0.4.0
#> [201] memoise_2.0.1               GenomicAlignments_1.47.0
#> [203] cluster_2.1.8.1             shinyWidgets_0.9.0
#> [205] shinyAce_0.4.4
```
